# Establishment of snake venom gland organoids from a novel family, Colubridae

**DOI:** 10.64898/2026.03.27.714740

**Authors:** Stephanie French, Rachael Da Silva, Rohit Patel, Claire H Caygill, Shannon Quek, Adam Westhorpe, Jens Puschhof, Rebecca Edge, Charlotte Dawson, Edouard Crittenden, Paul Rowley, Zachary Holland, Stephen P Mackessy, Cassandra M. Modahl

## Abstract

Non-front-fanged snakes are abundant, diverse and represent approximately 70% of extant snakes. However, there is limited knowledge about most species and their venoms, in part due to the technical and welfare challenges associated with venom extraction, low venom yields, and the lack of cellular models available. Organoids represent an excellent opportunity to overcome these challenges. Here, we establish, for the first time, venom gland organoids from snakes of the Colubridae family and demonstrate the *in vitro* production of toxins.

## Introduction

Venomous snakes can broadly be divided into front-fanged (Elapidae, Viperidae and Atractaspididae) and non-front-fanged (Ahaetuliinae, Calamariidae, Colubrinae, Dipsadidae, Grayiidae, Natricidae, Pseudoxenodontidae, Sibynophis, and Scaphiodontophis) species (Figueroa et al., 2016; Pyron et al., 2013; Zaher et al., 2019). Front-fanged snakes are the most medically important snakes, possessing enlarged, hollow maxillary teeth (fangs) and a venom gland with an internal reservoir for venom storage (Mackessy, 2022). However, non-front-fanged snakes are more diverse and widespread, comprising approximately 70% of all extant snake species (Mackessy and Saviola, 2016).

Non-front-fanged snakes possess enlarged or grooved solid rear maxillary teeth and a Duvernoy’s venom gland (Mackessy, 2022). Superficially similar to the venom gland of front fanged snakes, the Duvernoy’s venom gland is also located in the temporal region of the upper jaw. However, lacking an internal reservoir, venom is stored intracellularly and exocytosed into secretory tubules that flow into luminal ducts. Unlike the venom glands of front-fanged snakes, the Duvernoy’s venom gland is usually not surrounded by musculature that pressurise the gland, and venom is more slowly secreted around the base of the fang during prolonged biting (Kardong and Lavin-Murcio, 1993).

Understanding snake venom production, function and composition has mostly been focussed on front-fanged snakes. In line with this, organoids – self-organising 3D structures that grow from stem cells and produce organ-specific cell types through lineage differentiation (Clevers, 2016) – have previously been established from the venom glands of Elapidae and Viperidae. Snake venom gland organoids have been shown to recapitulate the venom gland through the production of functionally active toxins (Post et al., 2020). Here, we have established Duvernoy’s venom glands organoids from a non-front-fanged snake, *Boiga dendrophila*, and compare these organoids to those from *Bitis arietans* (Viperidae) venom gland. *B. dendrophila*, commonly known as the Mangrove Catsnake, is a large non-front fanged snake that is widespread across southeast Asia. *B. dendrophila* venom is primarily composed of prey-specific three-finger toxins (3FTxs) (Dashevsky et al., 2018; Mackessy and Saviola, 2016; Modahl and Mackessy, 2019), such as Boigatoxin-A (Lumsden et al., 2005) and Denmotoxin B (Pawlak et al., 2006). This new organoid model represents an excellent opportunity to study the evolution of venom systems, venom production and further characterise venom glands and toxins from rear-fanged snakes.

## Materials and Methods

### Venom gland tissue and venom

*Bitis arietans* venom was sourced from the on-site herpetarium at the Liverpool School of Tropical Medicine (LSTM). The facility and its protocols for the husbandry of snakes are approved by the UK Home Office and the LSTM Animal Welfare and Ethical Review Boards. *Boiga dendrophila* venom was kindly donated by Stephen Mackessy, University of Colorado. *Boiga* venom was collected following an IACUC approved protocol (0902E-SM-MBirdsL-12) as detailed by Hill & Mackessy with the use of 30 µg/g ketamine and 6.0 µg/g pilocarpine (Hill and Mackessy, 1997). All venoms were collected either prior to or in accordance with the Nagoya protocol.

Snake venom glands used in this study, specifically *Bitis arietans* (captive-bred and Nigerian locality; n=2) and *Boiga dendrophila* (captive-bred, n=2), came from snakes in the LSTM herpetarium. Snakes were euthanised by overdose of anaesthesia and cervical dislocation. Snake venom glands were dissected out as described in Puschhof *et al*. (Puschhof et al., 2021). Snake venom gland organoids from *Naja haje* (captive-bred) were supplied by Jens Puschhof and Yorick Post, Hubrecht Institute

### Organoid establishment

Organoids were setup using methods similar to those detailed in Puschhof *et al*. (2021). In brief, venom glands were either dissected from freshly euthanized snakes or collected from tissue stored in liquid nitrogen in 90% (v/v) fetal bovine serum, 10% (v/v) DMSO. Under sterile conditions, connective tissue was removed and gland cut into small pieces. Tissue was treated using am antibiotic cocktail media (DMEM, 50 µg/ml gentamicin, 2.5 µg/ml ciprofloxacin, 20 µM erythromycin, 10 nM azithromycin dehydrate, 100 µg/ml primocin) and digested using collagenase media solution (Advanced DMEM, 1 mg/ml collagenase, 10 µM Y-27632) at 32 °C, 5% CO_2_ for up to 1 h. Digested structures were plated at a density of approximately 100 per 15 µl in Cultrex basement membrane extract growth factor reduced type 2 (R&D Systems, 3533-001-02) and overlayed with expansion media (Advanced DMEM, 1x GlutaMAX, 1x HEPES, 100 U/ml penicillin-streptomycin, 100 µg/ml Primocin, 1.25 mM N-acetyl-L-cysteine, 1x B27 supplement, 10 mM nicotinamide, 1 µM prostaglandin E2, 2% (v/v) Noggin-conditioned media (IPA Therapeutics N002), 2% (v/v) R-spondin-conditioned media (IPA Therapeutics R002), 50 ng/ml human epidermal growth factor, 0.5 µM A83-91, 10 nM human gastrin-1, 100 ng/ml human fibroblast growth factor-10) containing 10 µM Y-27632, 50 µg/ml gentamicin, 2.5 µg/ml ciprofloxacin, 20 µM erythromycin and 10 nM azithromycin dehydrate. Expansion media was changed every other day, and organoids were passaged by mechanical dissociation with a 27G needle after 7 to 14 days.

### Organoid differentiation and lysate harvest

Organoids were passaged at least once prior to differentiation. Organoids were grown in expansion medium for 3 days then overlayed with differentiation medium (advanced DMEM, 1x GlutaMAX, 1x HEPES, 100 U/ml penicillin-streptomycin, 100 µg/ml Primocin, 1.25 mM N-acetyl-L-cysteine, 1x B27 supplement, 10 mM nicotinamide, 1 µM prostaglandin E2) for 7 days with media changed every other day. Organoid lysates were then harvested by sonication for 10s and protein concentration measured by the Qubit Protein Assay (Thermofisher Scientific) following manufactures instructions.

### Organoid staining

Organoids were harvested with Cultrex Organoid Harvesting Solution (R&D Systems, 3700-100-01). Active caspase-3 activity was determined using CellEvent Caspase-3/7 Gren detection reagent (Thermofisher Scientific, C10423). Organoids for hematoxylin and eosin (H&E) and periodic acid-Schiff (PAS) staining were resuspended in HistoGel (Fisher Scientific, 12006679) and staining outsourced to Liverpool Shared Research Facility, University of Liverpool.

### Western blot

Western blot was undertaken as detailed in Dawson *et al* with 1.6 μg of organoid lysates and 0.64 μg venom (Dawson et al., 2024). Antivenom used for the detection of *B. arietans* toxins was EchiTAb-Plus-ICP (lot 7131123PALF, Expiry: 11/2028), a polyspecific antivenom raised against a mixture of African snakes including *B. arietans*. Neuro Polyvalent Snake Antivenom (NPAV, lot NP00515, Expiry: 29/9/2020) – an antivenom targeting neurotoxic venoms from *Naja kaouthia, Ophiophagus hannah, Bungarus candidus* and *B. fasciatus* – was used for the detection of *B. dendrophila* toxins. Antivenoms were standardised to 10mg/ml and then diluted 1: 1,000. As in Dawson *et al*., Horse IgG (H&L) Antibody DyLight™800 Conjugated (Cambridge Bioscience, 608445002, 1:15,000) was utilised as the secondary antibody.

### Activity assays

Organoids protein lysates (200 µg/ml) and venom (100 µg/ml – 10,000 µg/ml) were analysed for snake venom metalloproteinase (SVMP) and phospholipase A_2_ (PLA_2_) activity. SVMP activity was determined *in vitro* using a quenched fluorogenic substrate (ES010 Mca-KPLGL-Dpa-AR-NH2, R&D Bio-systems) as detailed previously (Menzies et al., 2022). PLA_2_ activity was determined *in vitro* using the secretory phospholipase A_2_ assay kit from Abcam (AB133089) following manufacture instructions. Inhibition of muscle-type nicotinic acetylcholine receptor (nAChR) by *B. dendrophila* organoid protein lysates (60 µg/ml) and venom (0.1 µg/ml - 600 µg/ml) was determined using a membrane potential dye on TE671 cells expressing the γ-subunit containing muscle-type nAChR, as detailed in Patel *et al*. (Patel et al., 2023). Alpha-bungarotoxin (α-BgTx), isolated from *Bungarus multicinctus* venom was purchased from Invitrogen (B1601) and used as a positive control at 90nM in the nAChR inhibition assay.

### Statistical analysis

Data visualisation and statistical analysis was undertaken GraphPad Prism 10.2.1. Normality was assessed by Shapiro-Wilk test and visualisation with Q-Q plots, and parametric or non-parametric statistical tests were undertaken as appropriate. Differences in SVMP activity were assessed by comparing area under the curve. Differences in PLA_2_ activity were assessed by comparing the final time points (60 min). Differences in inhibition of activated nAChR were assessed by normalization to control acetylcholine (ACh)-induced nAChR activation.

## Results and Discussion

### Establishment of Colubridae organoids

*Boiga dendrophila* venom gland was digested, resulting in the release of small structures (Figure 1A, day 0) that generated proliferating organoids (Figure 1A, day 2 - 7). For comparisons to front-fanged snakes, *Bitis arietans* venom glands organoids were also established. Digestion of *B. arietans* venom glands resulted in the release of structures larger than that of *B. dendrophila* (Figure 1A, day 0) which formed organoids that expanded over time (Figure 1A, day 2 - 7). Expansion of both *B. dendrophila* and *B. arietans* organoids was challenging, and therefore *Naja haje* (Elapidae) organoids were cultured as a control (Figure 1A). These organoids were non-granular and expanded rapidly. In accordance, the only published accounts of long-term culture of snake venom gland organoids are those established from Elapidae (Mackessy, 2022; Puschhof et al., 2021).

**Figure 1.**
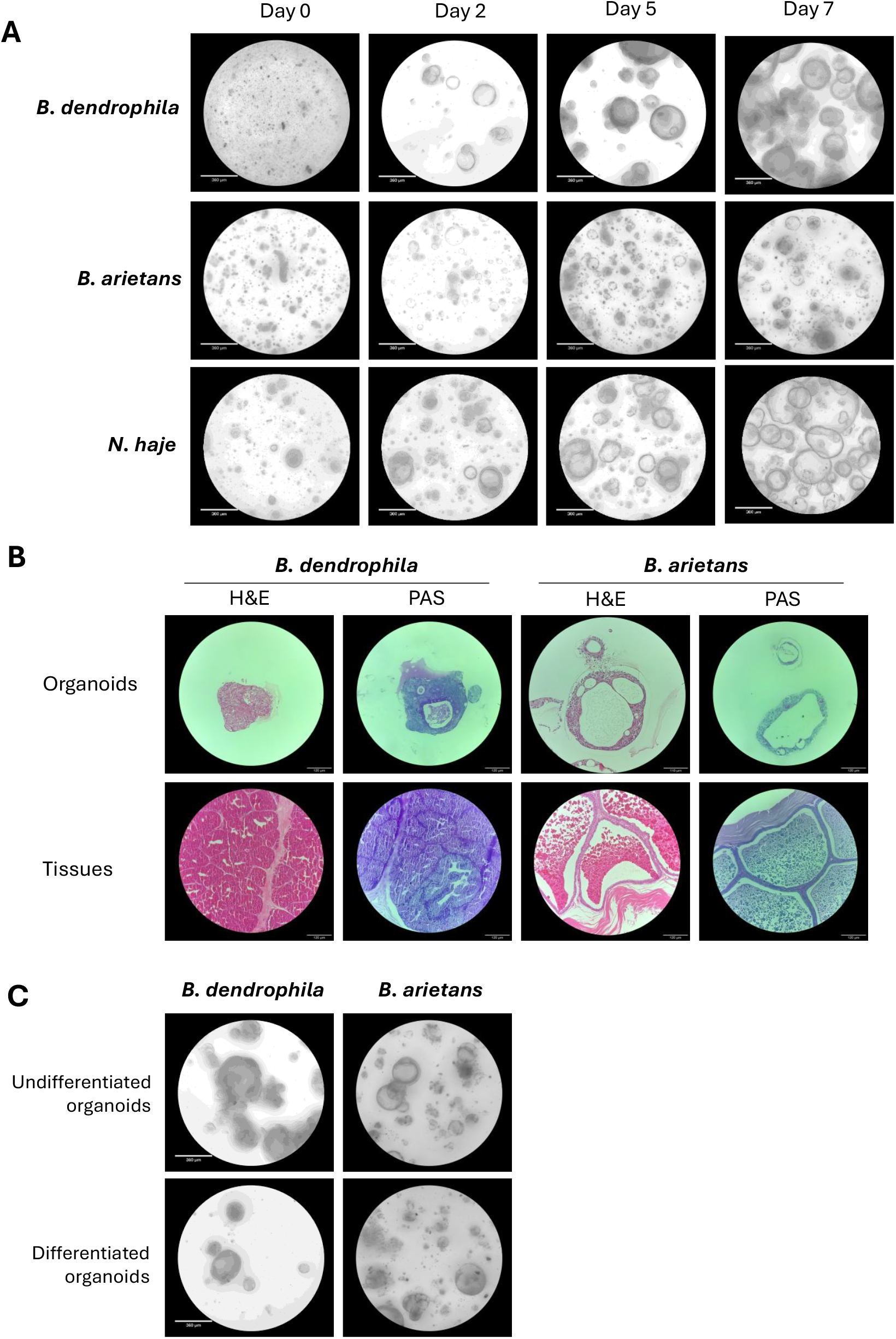
Establishment, growth and differentiation of snake venom gland organoids. Tissue fragments were isolated from freshly excised tissue of snake venom gland of *Boiga dendrophila* and *Bitis arietans* and expanded for 7 days in proliferation media. *Naja haje* organoids were revived from liquid nitrogen and expanded. **A)** Bright field images were taken to monitor organoid formation and growth. **B)** *B. dendrophila* and *B. arietans* organoids were stained with H&E and PAS. **C)** Light microscopy of differentiated and undifferentiated *B. dendrophila* and *B. arietans* venom gland organoids.

We then assessed whether *B. dendrophila* organoids could be established from tissue stored in liquid nitrogen, because freshly excised snake venom gland tissue is not readily available to many researchers. Establishment of *B. dendrophila* organoids from stored tissue samples, utilising the same method as on fresh tissue, was unsuccessful (Supplementary Figure 1).

### Histology of Colubridae organoids

Organoid morphology and cellular organization was assessed by H&E and PAS staining and compared to native tissue. Organoids showed a high level of similarity to native tissue. The majority of *B. dendrophila* organoids were lobular in shape composed of thick swirls of cells around a small lumen (Figure 1B), with smaller cavities and thicker cellular linings than *B. arietans* organoids. Some *B. dendrophila* organoids had a more spherical morphology (Supplementary Figure 2). In organoids from both species, PAS-positive granules and cells in the luminal lining confirmed the presence of mucosecretory cells, akin to tubules in the tissue.

Further analysis, such as single cell RNA sequencing and comparisons of transmission electron micrographs, will be required to compare cell types (Mackessy, 2022).

Organoids were then differentiated, resulting in reduced organoid expansion. No visible changes in *B. dendrophila* organoid morphology were observed under light microscope (Figure 2C). However, differentiation of *B. arietans* organoids appeared to induce some cell death. This was confirmed by active caspase-3 staining (Supplementary Figure 3). In addition, some extracellular matrix domes were completely degraded upon differentiation of *B. arietans* organoids, resulting in organoids becoming two-dimensional (Supplementary Figure 4). It is tempting to speculate that this could be due to SVMPs degrading the basement membrane extract (BME). Consistent with this hypothesis, BME breakdown has been observed in cultures of *Crotalus atrox* (Viperidae) venom gland organoids, but not *Crotalus scutulatus* venom gland organoids, which are species with SVMP-rich and SVMP-poor venom, respectively (Dr Timothy Peterson, University of Maryland, personal communications).

**Figure 2.**
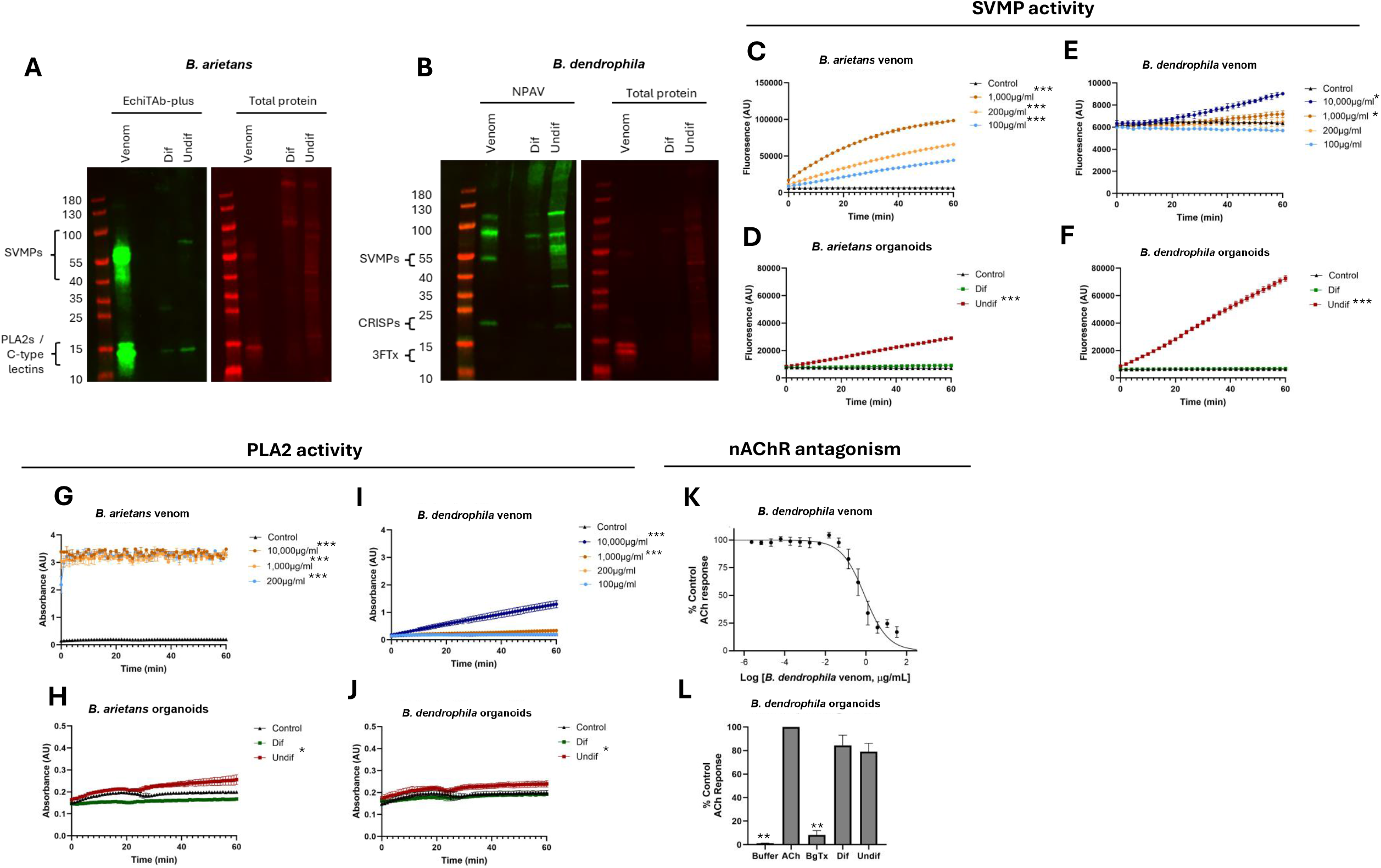
Toxin composition and activity of *B. dendrophila* and *B. arietans* venom and venom gland organoids. **A)** Toxin composition of *B. arietans* organoids and venom were analysed by Western blot using antivenom EchiTAb-plus. **B)** Toxin composition of *B. dendrophila* organoids and venom were analysed by Western blot using antivenom NPAV. **C-F)** SVMP activity of *B. arietans* venom (C), *B. arietans* organoids (D), *B. dendrophila* venom (E), *B. dendrophila* organoids (F). **G-J)** PLA2 activity of *B. arietans* venom (G), *B. arietans* organoids (H) *B. dendrophila* venom (I) and *B. dendrophila* organoids (J). **K-L)** Nicotinic acetylcholine receptor inhibition by *B. dendrophila* venom (K) and organoids (L). * p< 0.05, ** p< 0.05, *** p<0.001 relative to control for SVMP and PLA2 assay, and relative to ACh for neurotoxicity assay. Abbreviations: 3FTx, three finger toxins; ACh, acetylcholine; BgTx, Bungarotoxin; CRISPs, cysteine-rich secretory proteins; Dif, differentiated organoids; PLA2, phospholipase 2; SVMP, snake venom metalloproteinases; Undif, undifferentiated organoids,

### Venom toxin content in organoids lysates

Next it was assessed whether *B. dendrophila* and *B. arietans* venom gland organoids produced venom similar in composition to living snakes (Figure 2A-B). Western blotting of *B. arietans* venom revealed EchiTAb-Plus-ICP immunoreactive proteins at approximately 10 - 15 kDa and 40 - 70 kDa, consistent with the molecular weights of SVMPs and PLA_2_s (Bartlett et al., 2025; Howard, 1975). The low molecular weight band (10 - 15k Da) was observed in both differentiated and undifferentiated *B. arietans* organoid lysates, whilst a band at ~70 kDa was only observed in undifferentiated organoids. These findings were also observed in organoids lysates from *B. arietans* with Nigerian locality (Supplementary Figure 5), suggesting the production of SVMPs and PLA_2_s in undifferentiated *B. arietans* organoids.

Due to the lack of *Boiga* spp. antivenom, Neuro Polyvalent Snake Antivenom (NPAV) – an antivenom raised against Asian species with high 3FTx content – was utilised in an attempt to detect *B. dendrophila* toxins. NPAV had cross reactivity with *B. dendrophila* venom, but only low immunoreactivity towards 3FTxs (10 - 15 kDa) (Supplementary Figure 6). Immunoreactive proteins at 15 - 25 kDa, 55 kDa, 55 - 100 kDa and 100 - 130 kDa were seen for the undifferentiated organoids and *B. dendrophila* venom. Bands at 55 to 100 kDa were detected in differentiated *B. dendrophila* organoids (Figure 2A). Using information from published accounts of *Boiga* venom composition, we speculate the 20 kDa organoid toxins detected to be CRISPs (Dashevsky et al., 2018) and the ~55 kDa toxin to be SVMPs (Pla et al., 2018). The two toxins in *dendrohila* venom at 55 - 100 and 100 - 130 kDa have been detected previously but remain uncharacterised (Pla et al., 2018), preventing us from identifying these toxins in the organoids. We were unable to confirm the presence of 3FTxs (toxins smaller than 15 kDa, typically 6 – 8.1 kDa) in organoids, potentially due to the low reactivity NPAV against *B. dendrophila* 3FTxs. These findings demonstrate the capacity of non-front fanged snake venom gland organoids to produce toxins that reflect the protein composition of whole venom. However, top-down proteomics and transcriptomics analysis will be required to validate the presence of specific toxins and further understand the differences between differentiated and undifferentiated venom gland organoids.

Total protein staining showed the presence of many additional bands in the organoid lysates that were not present in venom collected from either *B. arietans or B. dendrophila*. This is because venom was harvested from organoids by sonication, resulting in the release of intracellular non-toxin proteins. It would be interesting to explore alternative methods of organoid toxin harvest by mechanical disruption such as freeze-thaw and repeat pipetting with a narrow tip, in an attempt to reduce non-toxin protein content.

### Venom toxin activity in organoids lysates

Next, we assessed if the toxins produced by venom gland organoids were functional. In accordance with the SVMP activity detected in *B. arietans* venom (100 - 1,000 µg/ml, ANOVA, p < 0.001, Figure 2C) and *B. dendrophila* venom (1,000 µg/ml - 10,000 µg/ml, ANOVA, p < 0.05, Figure 2D), undifferentiated *B. arietans* and *B. dendrophila* organoid lysates displayed significant SVMP activity (ANOVA, p < 0.001, Figure 2E-F). Differentiated organoid lysates did not have SVMP activity (ANOVA, p >> 0.05, Figure 2E-F), consistent with the lack of antivenom-reactive proteins at ~55kDa in these samples.

In accordance with previous studies (Broaders and Ryan, 1997; Hill and Mackessy, 2000; Howard, 1975), PLA_2_ activity was observed in *B. arietans* venom (100 - 1,000 µg/ml, ANOVA, p < 0.001, Figure 2G) and *B. dendrophila* venom (1,000 - 10,000 µg/ml, ANOVA, p < 0.05, Figure 2H). Again, undifferentiated organoid lysates had PLA_2_ activity (ANOVA, p < 0.05, Figure 2I-J), whilst differentiated organoid lysates lacked activity (ANOVA, p > 0.05, Figure 2I-J).

*B. dendrophila* venom displayed dose-dependent α-neurotoxin activity; inhibiting ACh-induced activation of the muscle-type nicotinic acetylcholine receptor (nAChR) with an IC_50_ of 0.86 µg/ml (Figure 2K). Interestingly, this is in the same order of magnitude as *Dendroaspis viridis* venom (IC50 of 0.59 µg/ml) (Patel et al., 2023) and represent the first report of *B. dendrophila* venom or toxin inhibiting a human nAChR. Previous studies have examined the chick or mouse nAChR (Lumsden et al., 2004; Lumsden et al., 2005; Pawlak et al., 2006). Dosing with differentiated and undifferentiated organoids slightly decreased ACh-induced nAChR activation, however, this was not significant (ANOVA, p > 0.05, Figure 2L). This is potentially due to the lower concentration used compared to the PLA_2_ and SVMP assay, as samples had to be diluted to obtain the larger volume required for this assay. It would have been interesting to repeat experiments with higher concentrations of organoid lysates. However, the limited expansion of the organoids prevented this and highlights the need for further studies optimising the sub-culture of this family of organoids.

These findings hint towards the production of functionally active toxins in organoids, however, it is important to note the presence of intracellular non-toxin proteins in organoid lysates that could interfere with assay results. It will be important to isolate and test specific organoid toxins by HPLC to test this hypothesis.

In conclusion, this is the first establishment of snake venom gland organoids from non-front-fanged snakes. Venom extraction from this group of snakes is challenging, providing very limited yields and requiring extraction chemicals which contaminate the venom (Damm et al., 2025). This research takes a step towards utilising organoids as a tractable and renewable platform for venom production and functional studies. *Boiga spp*. organoids specifically are a valuable model to investigate evolutionary adaptations due to their prey-specific toxins, and serve as a comparative baseline for the more specialised venom glands of front-fanged snakes. Non-front-fanged organoids provide a powerful platform to investigate many aspects of snake venom glands, from establishing cell types and lineages, to identifying drivers of venom diversity, to studying venom synthesis and secretion (Mackessy, 2022).

## Abbreviations

3FTx: three-finger toxins
BME: basement membrane extract
Ach: acetylcholine
BSA: bovine serum albumin
DMEM: Dulbecco’s Modified Eagle’s Medium
DMSO: dimethyl sulfoxide
FBS: fetal calf serum
H&E: hematoxylin and eosin
LSTM: Liverpool School of Tropical Medicine
PAS: periodic acid-Schiff
PBS: phosphate buffered saline
PLA_2_: phospholipase A2
nAChR: nicotinic acetylcholine receptor
SVMP: snake venom metalloproteinases

## Acknowledgements

Thank you to Gemma Charlesworth and Marie O’Brien at the University of Liverpool Shared Research Histology Core Facility for undertaking the H&E and PAS staining (RRID:SCR_026606). Thank you to Grant Hughes and Shaun Pennington for instrument support.

## References

Bartlett, K. E., Westhorpe, A., Wilkinson, M. C. and Casewell, N. R. (2025). Snake venom metalloproteinases from puff adder and saw-scaled viper venoms cause cytotoxic effects in human keratinocytes. Toxins 17, 328.

Broaders, M. and Ryan, M. F. (1997). Enzymatic properties of the Duvernoy’s secretion of Blanding’s Tree snake (Boiga blanding) and the Mangrove Snake ((Boiga dendrophila). Toxicon 35, 1148.

Clevers, H. (2016). Modeling development and disease with organoids. Cell 165, 1586–1597.

Damm, K., Hartwig, C., Vilcinskas, A., Mackessy, S. P., Lüddecke, T. and Damm, M. (2025). Ketamine and metabolites in snake venom: effects of venom extraction and potential impact on animal models. Sci. Rep. 15, 1–5.

Dashevsky, D., Debono, J., Rokyta, D., Nouwens, A., Josh, P. and Fry, B. G. (2018). Three-finger toxin diversification in the venoms of cat-eye snakes (Colubridae: Boiga). J. Mol. Evol. 86, 531–545.

Dawson, C. A., Bartlett, K. E., Wilkinson, M. C., Ainsworth, S., Albulescu, L. O., Kazandijan, T., Hall, S. R., Westhorpe, A., Clare, R., Wagstaff, S., et al. (2024). Intraspecific venom variation in the medically important puff adder (Bitis arietans): Comparative venom gland transcriptomics, in vitro venom activity and immunological recognition by antivenom. PLoS Negl. Trop. Dis. 18, 1–22.

Figueroa, A., McKelvy, A. D., Grismer, L. L., Bell, C. D. and Lailvaux, S. P. (2016). A species-level phylogeny of extant snakes with description of a new colubrid subfamily and genus. PLoS One 11, 1–31.

Hill, R. E. and Mackessy, S. P. (1997). Venom yields from several species of colubrid snakes and differential effects of ketamine. Toxicon 35, 671–678.

Hill, R. E. and Mackessy, S. P. (2000). Characterization of venom (Duvernoy’s secretion) from twelve species of colubrid snakes and partial sequence of four venom proteins. Toxicon 38, 1663–1687.

Howard, N. L. (1975). Phospholipase As from puff adder (Bitis arietans) venom. Toxicon 13, 21–30.

Kardong, K. V. and Lavin-Murcio, P. A. (1993). Venom delivery of snakes as high-pressure and low-pressure systems. Copeia 1993, 644–650.

Lumsden, N. G., Fry, B. G., Manjunatha Kini, R. and Hodgson, W. C. (2004). In vitro neuromuscular activity of ‘colubrid’ venoms: clinical and evolutionary implications. Toxicon 43, 819–827.

Lumsden, N. G., Fry, B. G., Ventura, S., Manjunatha Kini, R. and Hodgson, W. C. (2005). Pharmacological characterisation of a neurotoxin from the venom of Boiga dendrophila (Mangrove catsnake). Toxicon 45, 329–334.

Mackessy, S. P. (2022). Venom production and secretion in reptiles. Journal of Experimental Biology 225, 1–10.

Mackessy, S. P. and Saviola, A. J. (2016). Understanding biological roles of venoms among the Caenophidia: the importance of rear-fanged snakes. Integr. Comp. Biol. 56, 1004–1021.

Menzies, S. K., Clare, R. H., Xie, C., Westhorpe, A., Hall, S. R., Edge, R. J., Alsolaiss, J., Crittenden, E., Marriott, A. E., Harrison, R. A., et al. (2022). In vitro and in vivo preclinical venom inhibition assays identify metalloproteinase inhibiting drugs as potential future treatments for snakebite envenoming by Dispholidus typus. Toxicon X 14, 1–10.

Modahl, C. M. and Mackessy, S. P. (2019). Venoms of rear-fanged snakes: new proteins and novel activities. Front. Ecol. Evol. 7, 1–18.

Patel, R. N., Clare, R. H., Ledsgaard, L., Nys, M., Kool, J., Laustsen, A. H., Ulens, C. and Casewell, N. R. (2023). An in vitro assay to investigate venom neurotoxin activity on muscle-type nicotinic acetylcholine receptor activation and for the discovery of toxin-inhibitory molecules. Biochem. Pharmacol. 216, 1–13.

Pawlak, J., Mackessy, S. P., Fry, B. G., Bhatia, M., Mourier, G., Fruchart-Gaillard, C., Servent, D., Ménez, R., Stura, E., Ménez, A., et al. (2006). Denmotoxin, a three-finger toxin from the colubrid snake Boiga dendrophila (mangrove catsnake) with bird-specific activity. Journal of Biological Chemistry 281, 29030–29041.

Pla, D., Petras, D., Saviola, A. J., Modahl, C. M., Sanz, L., Pérez, A., Juárez, E., Frietze, S., Dorrestein, P. C., Mackessy, S. P., et al. (2018). Transcriptomics-guided bottom-up and top-down venomics of neonate and adult specimens of the arboreal rear-fanged Brown Treesnake, Boiga irregularis, from Guam. J. Proteomics 174, 71–84.

Post, Y., Puschhof, J., Beumer, J., Kerkkamp, H. M., de Bakker, M. A. G., Slagboom, J., de Barbanson, B., Wevers, N. R., Spijkers, X. M., Olivier, T., et al. (2020). Snake venom gland organoids. Cell 180, 233-247.e21.

Puschhof, J., Post, Y., Beumer, J., Kerkkamp, H. M., Bittenbinder, M., Vonk, F. J., Casewell, N. R., Richardson, M. K. and Clevers, H. (2021). Derivation of snake venom gland organoids for in vitro venom production. Nat. Protoc. 16, 1494–1510.

Pyron, R. A., Burbrink, F. T. and Wiens, J. J. (2013). A phylogeny and revised classification of Squamata, including 4161 species of lizards and snakes. BMC Evol. Biol. 13, 1–53.

Zaher, H., Murphy, R. W., Arredondo, J. C., Graboski, R., Machado-Filho, P. R., Mahlow, K., Montingelli, G. G., Quadros, A. B., Orlov, N. L., Wilkinson, M., et al. (2019). Large-scale molecular phylogeny, morphology, divergence-time estimation, and the fossil record of advanced caenophidian snakes (Squamata: Serpentes). PLoS One 14, 1–82.

