## Supplementary Figures 1-6 for "Establishment of snake venom gland organoids from a novel family, Colubridae"

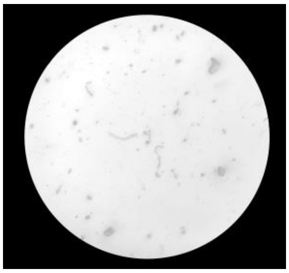

**Supplemental Figure 1. Unsuccessful establishment of *Boiga dendrophila* snake venom gland organoids from cryopreserved tissue.** Bright field images were taken at day 7 to assess organoid formation and growth.

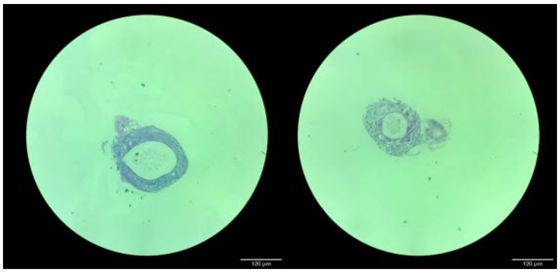

**Supplementary Figure 2. Additional PAS images of undifferentiated *Boiga dendrophila* organoids with spherical morphology.**

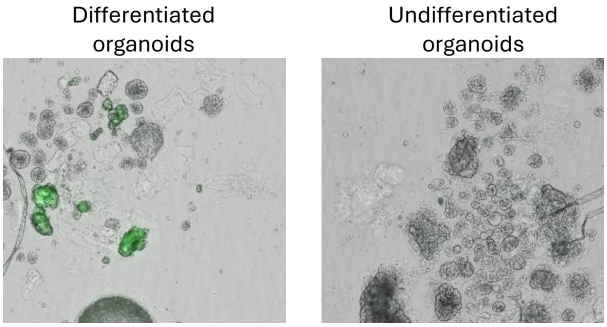


**Supplemental Figure 3. Active caspase 3 staining of *Bitis arietans* organoids.** Differentiated organoids and undifferentiated organoids were incubated with CellEvent Caspase- 3/7 Gren detection reagent to detect active caspase 3 (green). Bright field images and fluorescence images were taken.

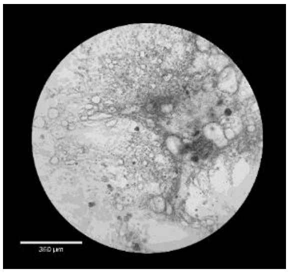

**Supplemental Figure 4. Extracellular matrix degradation by *Bitis arietans* snake venom gland organoids.** Bright field images at day 7 of differentiation.

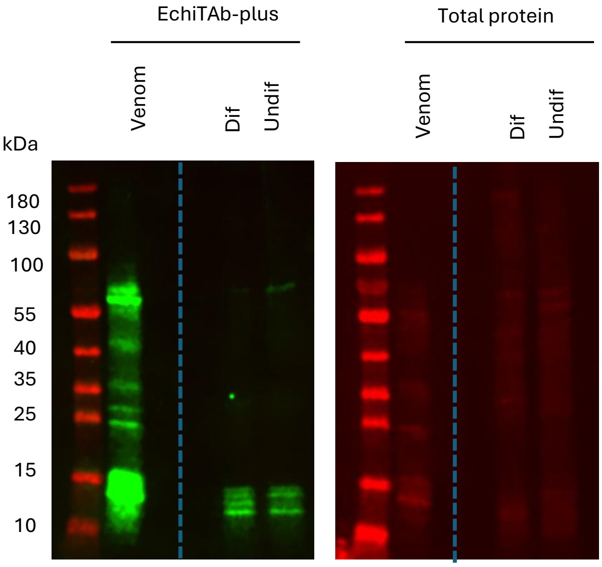


**Supplementary Figure 5. Toxin composition *Bitis arietans* venom gland organoids lysates with Nigerian locality.** Toxin composition of *B. arietans* venom, differentiated organoids (Dif) and undifferentiated organoids (Undif) were analysed by Western blot.

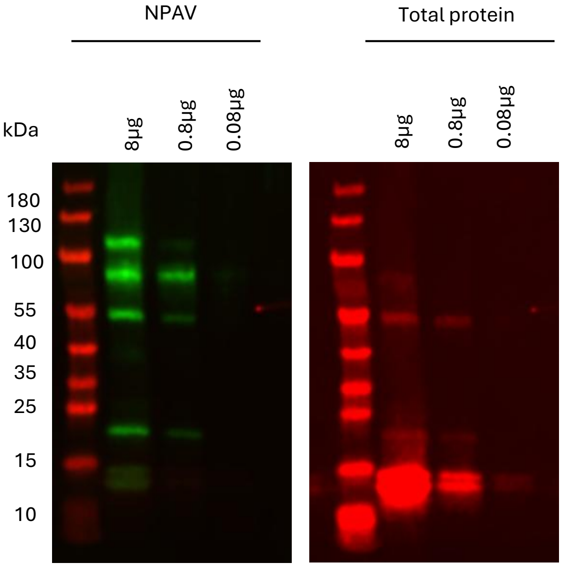


**Supplementary Figure 6. Confirmation of NPAV antivenom detecting *B. dendrophila* venom toxins.** Binding of NPAV to *B. dendrophila* venom was assessed with 3 different protein contents (8μg, 0.8μg, 0.08μg).
